# Automatic Auditory Streaming Restores Missing Temporal Modulations in Echoic Speech

**DOI:** 10.1101/2023.05.11.540309

**Authors:** Jiaxin Gao, Mingxuan Fang, Honghua Chen, Nai Ding

**Affiliations:** Key Laboratory for Biomedical Engineering of Ministry of Education, College of Biomedical Engineering and Instrument Sciences, Zhejiang University, Hangzhou 310027, China; The State key Lab of Brain-Machine Intelligence; The MOE Frontier Science Center for Brain Science & Brain-machine Integration, Zhejiang University, Hangzhou, 310027, China

## Abstract

Human listeners can reliably recognize speech in adverse listening environments, and previous studies have identified that reliable neural encoding of slow temporal modulations in speech is essential for speech recognition. Recent behavioral studies demonstrate that long-delay echoes, which are rare in physical environments but common during online conferencing, can eliminate critical temporal modulations. These echoes, however, barely affect speech intelligibility, and here we investigate the underlying neural mechanisms. MEG experiments demonstrate that cortical activity can effectively track the temporal modulations eliminated by an echo, which cannot be explained by basic neural adaptation mechanisms such as synaptic depression, gain control, and adaptive filtering. Instead, the cortical response to echoic speech is better explained by a model that segregates speech from its echo than a model that encodes echoic speech as a whole. The speech segregation effect is observed even when attention is diverted, but disappears when speech segregation cues in the spectro-temporal fine structure are degraded. Altogether, these results strongly suggest that the auditory system can automatically segregate speech and its echo and encode them as two auditory streams, providing a potential neural basis for reliable speech recognition in echoic environments.

## Introduction

To reliably extract auditory objects from a complex auditory scene is a primary goal of auditory processing^1–4^. For humans, speech is the most important communicational sound and humans can reliably recognize speech in various listening environments^5, 6^. Previous studies have suggested that the neural representation of speech envelope, i.e., the fluctuation in speech power over time^7, 8^, is resilient to many types of acoustic degradation at the level of auditory cortex^9–15^. Previous studies have shown that distinct neural mechanisms are required to compensate different types of acoustic degradation. For example, stationary noise can greatly attenuate the dynamic range of temporal modulations^16^, an effect that can be well compensated by relatively basic neural adaptation mechanisms^9, 10, 17^. When speech is mixed with a competing voice, however, neural adaptation to the overall statistical properties of the sound mixture can barely help to enhance the neural response to the target speech stream. Instead, the brain has to segregate the sound mixture into auditory streams by analyzing, e.g., pitch and spatial cues, and selectively process the auditory stream of interest under the modulation of top-down attention^2–4, 18^. In such conditions, if speech segregation cues are removed through, e.g., noise vocoding, speech segregation fails and the speech recognition rate drops^19, 20^.

A large number of studies have focused on the neural encoding of speech envelope, since the speech envelope provides crucial cues for speech recognition^21, 22^. When the speech envelope is further decomposed into faster and slower components, referred to as temporal modulations^23, 24^, it has been demonstrated that temporal modulations between 1 and 8 Hz are essential for speech recognition: When the temporal modulations between 1 and 8 Hz are removed, a large number of studies consistently reported that speech intelligibility drops^25–28^. Recently, however, it is demonstrated that a single echo can eliminate the temporal modulations at frequencies that are determined by the echo delay. For example, an echo with 125-ms delay can eliminate temporal modulations at 4 Hz, and is supposed to reduce speech intelligibility. Nevertheless, young listeners show no difficulty recognizing such echoic speech^29^. The high intelligibility of echoic speech indicates that either the temporal modulations eliminated by an echo are neurally restored or that these temporal modulations are actually not necessary for speech recognition, in contradiction to the conclusions of a large number of previous studies^25–28^.

Echoes can be viewed as a special case for reverberation, since both echoes and reverberation are generated by sound reflections. For a single echo, the power of sound reflection concentrates at a single delay (Figure 1A, B). In daily reverberant environments, however, the power of sound reflection decays exponentially as the delay of the reflection increases^30^ (Figure 1B). Echoic environments are relatively rare in physical environments, but are prevalent during online conferencing^31^. Reverberations in daily environments^27^ create moderate frequency-dependent attenuation to the speech temporal modulations, while an intense echo can completely eliminate the temporal modulations at frequencies that are determined by the echo delay^29^ (Figure 1C). Previous studies have shown the influence of reverberation on speech envelope can be compensated through basic neural adaptation mechanisms^9, 32^. It remains unclear, however, whether the auditory system also has mechanisms to restore the temporal modulations that are completely eliminated by echoes.

**Figure 1:**
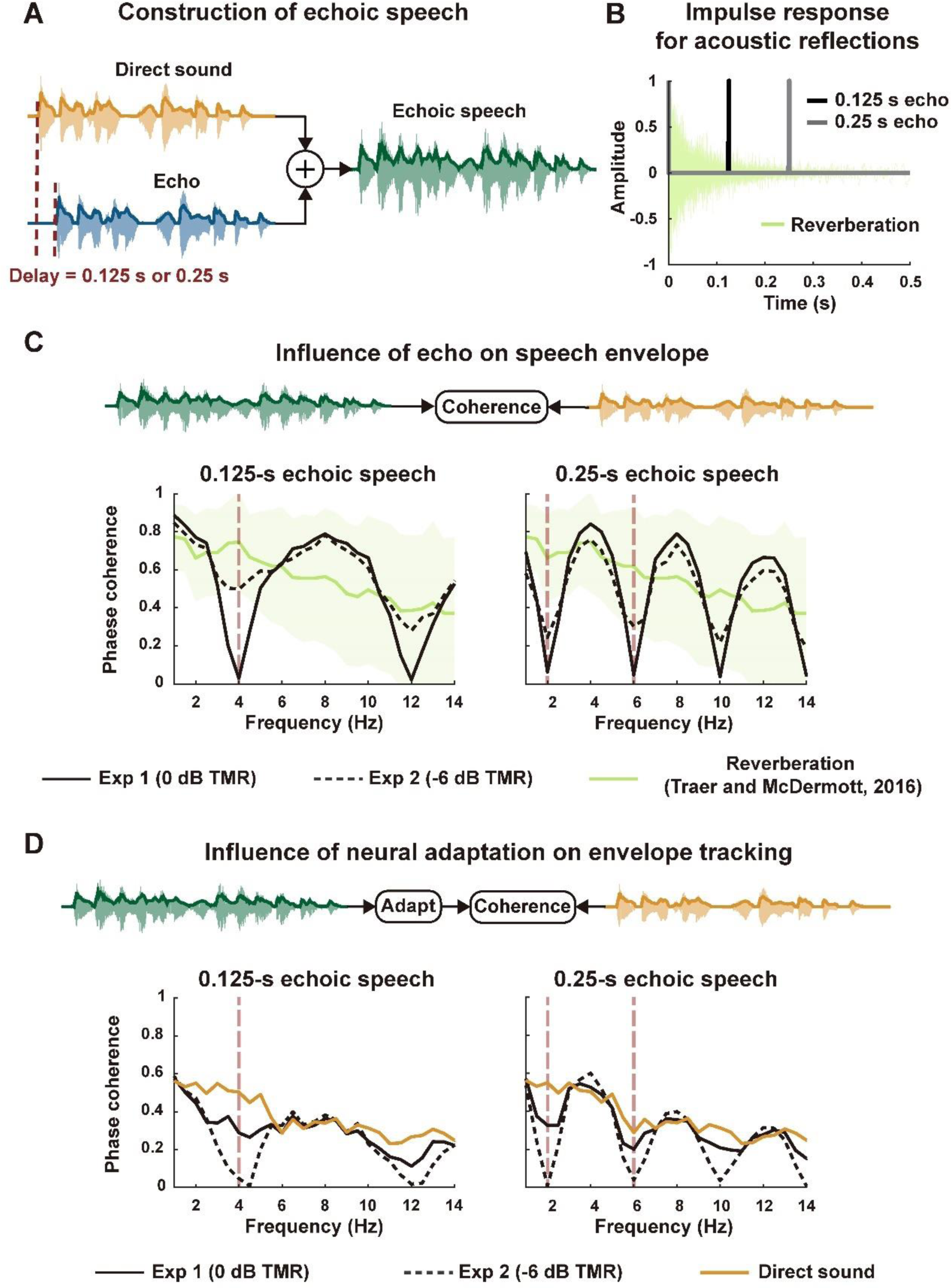
Properties of echoic speech and neural adaptation models. **A,** Echoic speech in Experiments 1 and 2, which is generated by mixing a direct sound with an echo, which are identical except for a time delay. The delay is either 0.125 s or 0.25 s. For the Experiment 1, the echo has the same amplitude as the direct sound (0 dB TMR). For the Experiment 2, the echo is twice stronger than the direct sound (−6 dB TMR). **B,** The impulse response of reverberant and echoic environments. The thin green curves show the impulse responses of 270 reverberant environments reported by Traer and McDermott (2016). The thick gray and black curves show the equivalent impulse response for acoustic reflections for the echoic speech analyzed in the current study. **C,** Influence of echo and reverberation on speech envelope, characterized by the phase coherence spectrum between the envelope of echoic/reverberant speech and the envelope of direct sound. Reverberant speech is constructed based on the 270 impulse responses reported by Traer and McDermott (2016), and the green curve and shaded area show the mean and the area between 2.5 and 97.5 percentiles. Dashed red line denotes the echo-related frequencies below 10 Hz. **D,** Phase coherence spectrum between the simulated neural response to echoic speech and the envelope of direct sound. Neural responses are simulated by processing the speech envelope using the neural adaptation model in Mesgarani et al. (2014) and the parameters are optimized to cancel the influence of echo. Since the neural adaptation model alters speech envelope, we also illustrate the result for anechoic speech, i.e., the direct sound alone. Analysis of the neural response simulated using another 2 models are shown in Figure S1. None of the models could effectively cancel the influence of echoes in all echoic conditions.

The current study investigates the neural encoding of echoic speech using MEG, and probes the underlying neural mechanisms through computational models. We added a single echo to a narrated story (Figure 1A), and manipulated the relative intensity and delay of echo to create strong influence on temporal modulations that are important to speech recognition. In such challenging echoic environments, we investigate using MEG whether auditory cortex can track the temporal modulations that are attenuated or eliminated by an echo. If neural activity can track these missing components, we further hypothesize that this is achieved through auditory stream segregation since an echo is subjectively perceived as a separate sound^33^. We test the hypothesis by analyzing whether speech and its echo have distinct spatio-temporal representations in the MEG response using the temporal response function (TRF) models.

## Results

### An echo severely distorts the speech envelope

Echoic speech was a mixture of two speech signals that only differed by a time delay (Figure 1a). In the following, the leading sound was referred to as the direct sound, and the lagging sound was referred to as the echo. We constructed challenging echoic conditions in which the echo was either of the same amplitude as the direct sound or was twice stronger. In other words, the target-to-masker ratio (TMR), i.e., the intensity ratio between the direct sound and the echo, was either 0 dB or −6 dB. The echo delay was either 0.125 s or 0.25 s (see methods for rationales).

We first characterized how an intense echo influenced the speech envelope at each modulation frequency using the phase coherence spectrum between the envelope of echoic speech and the envelope of the direct sound. The phase coherence spectrum calculated the phase locking between two signals in each narrow frequency band. The phase coherence value was 1 for two identical signals and was 0 for two independent signals. For speech with a 0.125-s echo, the phase coherence spectrum showed notches at 4 and 12 Hz. For speech with a the 0.25-s echo, notches were observed at 2, 6, 10, and 14 Hz (Figure 1C). In the following, frequencies corresponding to the notches in the phase coherence spectrum were referred to as the echo-related frequencies. Since previous studies showed that the cortical responses mainly track speech envelope below 10 Hz^34^, we only analyzed the echo-related frequencies below 10 Hz.

An intense echo severely distorted the speech envelope, but it was possible that the echo-related distortions were automatically compensated by the basic adaptation mechanisms in the auditory system. Here, we considered three types of neural computational models that were previously proposed to explain reliable envelope tracking in reverberant environments, i.e., synaptic depression, gain control, and linear adaptive filtering^9, 32, 35^. To evaluate whether these computations could lead to an echo-invariant speech representation, we calculated the phase coherence spectrum between the output of the models, i.e., the simulated neural responses, and the envelope of the direct sound (Figure 1D and Figure S1). All models predicted that neural tracking of the direct sound was attenuated at echo-related frequencies, compared to the neural tracking of anechoic speech.

### MEG response can track the speech envelope components notched out by an echo

Next, we analyzed how the actual speech-tracking neural response was influenced by an echo. Specifically, we calculated the phase coherence spectrum between the MEG response to echoic/anechoic speech and the envelope of the direct sound. We tested whether the neural tracking of the direct sound was attenuated at echo-related frequencies, as was predicted by the neural adaptation models. Experiments 1 and 2 separately tested echoic conditions in which the TMR was 0 dB or −6 dB, respectively. In Experiment 1, the participants listened to anechoic speech, echoic speech (0.125-s echo delay), and echoic speech (0.25-s echo delay) in separate blocks, and the cortical responses were recorded using MEG. The magnitude of the echo was the same as the magnitude of the direct sound (TMR = 0 dB). Participants were asked to attend to speech and answer comprehensions after each block. The mean question-answering accuracy was 95.6% in all three conditions. We characterized neural tracking of direct sound using the phase coherence spectrum between MEG responses and the temporal envelope of the direct sound. The phase coherence spectrum was calculated for each gradiometer and the average over all gradiometers was shown in Figure 2. In the anechoic condition, i.e., when the stimulus only included the direct sound, the phase coherence between MEG response and speech envelope was significantly above chance below ∼10 Hz (Figure 2A, *p* < 0.05, permutation test, FDR corrected).

**Figure 2:**
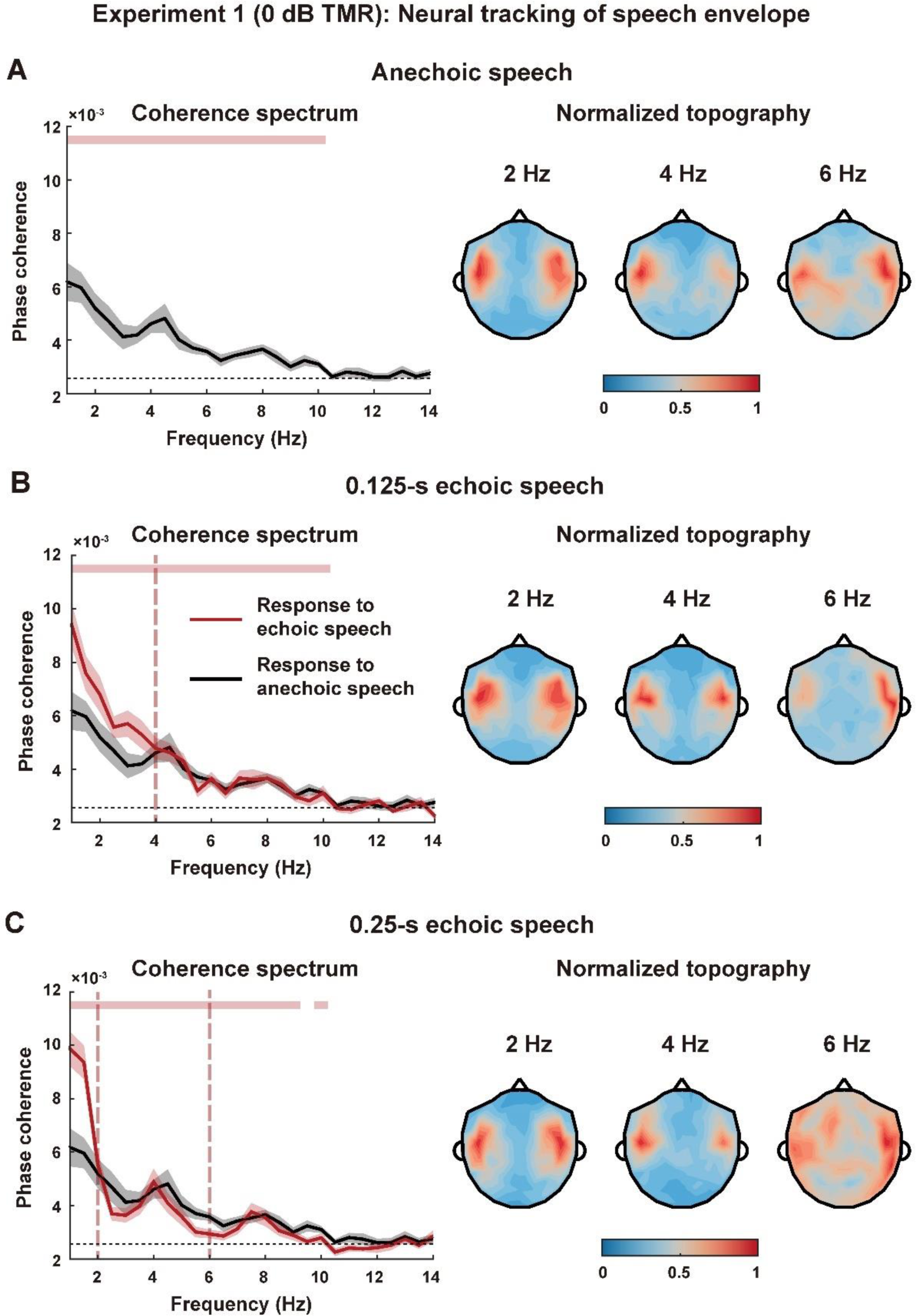
Results of Experiment 1. Panels **A, B,** and **C** show the results in the anechoic condition, 0.125-s echoic condition, and 0.25-s echoic condition, respectively. **Left:** the phase coherence spectrum between MEG response and the envelope of direct sound, which is averaged over participants and MEG gradiometers. The shaded areas cover 1 standard error across participants (*N* = 15) on each side. The dashed horizontal black line shows the chance-level phase coherence spectrum, and the dashed vertical red line denotes the echo-related frequencies. The pink bar on top denotes the frequency bins in which the phase coherence is significantly higher than chance (*p* < 0.05, permutation test, FDR corrected). **Right:** Topography of the phase coherence at echo-related frequencies. The topography only shows the results of gradiometers and the two gradiometers at the same position are averaged. Each topography is normalized by dividing its maximum.

For the echoic conditions, the phase coherence was also significantly above chance below ∼10 Hz (Figure 2B, C, *p* < 0.05, permutation test, FDR corrected), and no notches were observed at the echo-related frequencies (Figure 2B, C, *p* = 0.0002 for all echo-related frequencies, bootstrap, FDR corrected). The response topography at the echo-related frequency revealed bilateral temporal distribution (Figure 2). Compared with the anechoic condition, the 0.125-s echoic condition did not show a significant reduction in phase coherence at the echo-related frequency (Figure 3A, *p* = 0.91 at 4Hz, FDR corrected) and the 0.25-s echoic condition showed a significant reduction at 6 Hz (*p* = 0.0083, bootstrap, FDR corrected) but not at 2 Hz (*p* = 0.61 at 2Hz, bootstrap, FDR corrected). Furthermore, in contradiction to the prediction of neural adaptation models, we observed that the phase coherence around 1 Hz was enhanced in the echoic condition and this difference was statistically significant (Figure 3A, *p* = 0.0005 both in 0.125-s and 0.25-s echoic condition, bootstrap, FDR corrected). These results suggested that the human auditory system could effectively restore the speech envelope at the echo-related frequencies. The topography only shows the results of gradiometers and the two gradiometers at the same position are averaged. Each topography is normalized by dividing its maximum.

**Figure 3:**
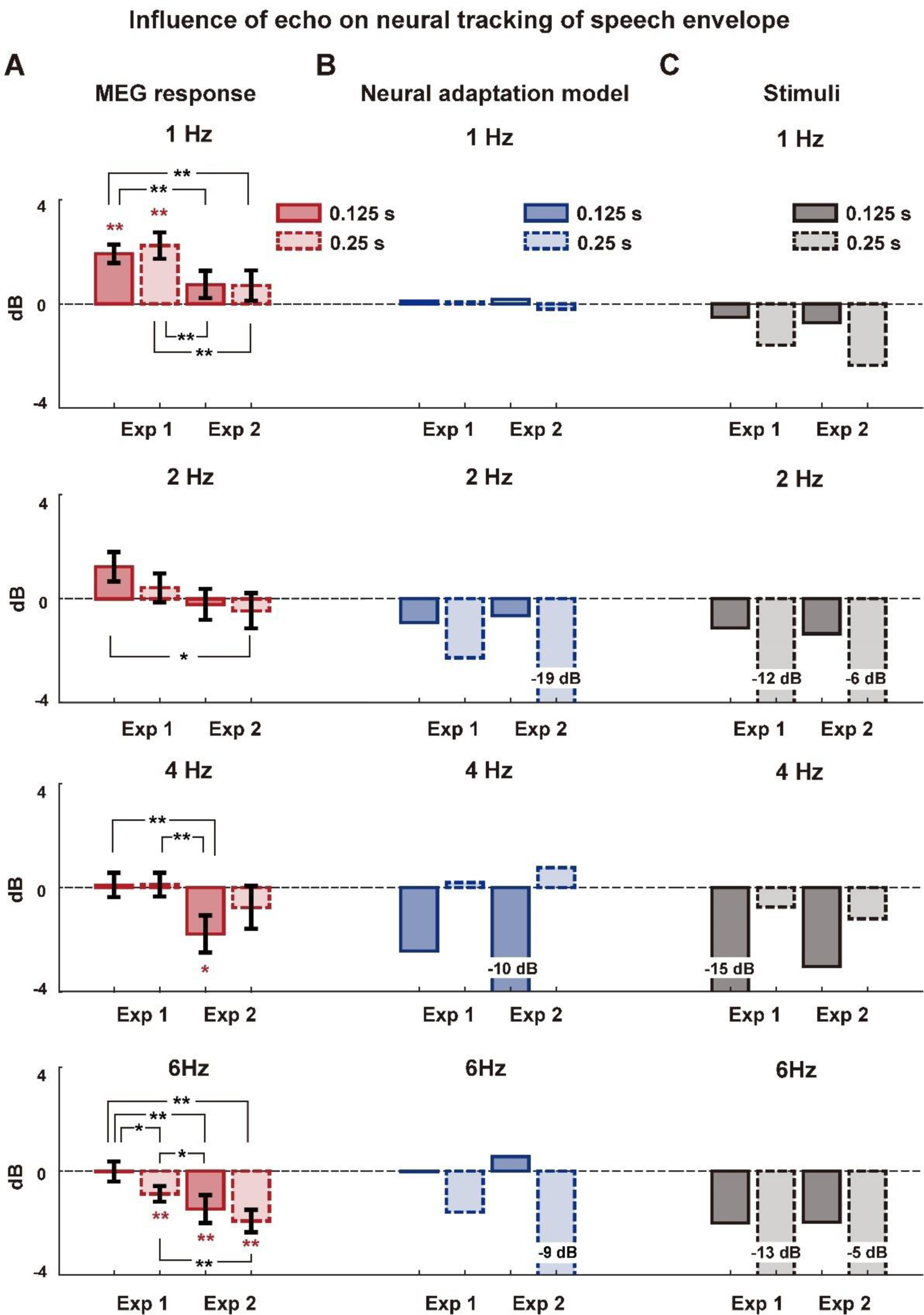
Influence of echo on the neural tracking of direct sound. The figure shows the difference in the MEG-envelope phase coherence between echoic and anechoic conditions, at echo-related frequencies. Specifically, for each participant, the phase coherence in the anechoic condition is subtracted from the phase coherence in the echoic condition. **A,** MEG results in Experiments 1 and 2. Red stars indicate that the difference in phase coherence significantly differs from 0. Black stars indicate a significant difference between two echoic conditions. (* *p* < 0.05, ** *p* < 0.01, bootstrap, FDR corrected). Error bars represent 1 standard error across participants (Experiment 1: *N* = 15; Experiment 2: *N* = 12). **B,** Results for the neural response simulated based on the neural adaptation model^9^ in Figure 1D. **C,** Results for the coherence between the envelope of echoic speech and the envelope of the direct sound.

Experiment 2 was the same as Experiment 1, except that the TMR was set to −6 dB. In other words, the echo was twice stronger than the direct sound. We tested this condition since simulations showed that neural adaptation models tended to aggravate the loss of speech envelope at echo-related frequencies (Figure 1D). Participants correctly answered 94.4%, 97.2%, and 100% questions in the anechoic, 0.125-s echoic and 0.25-s echoic conditions. In all three conditions, the phase coherence between MEG responses and the envelope of direct sound was above chance level below 10 Hz (Figure 4, *p* < 0.05, permutation test, FDR corrected), except for the 6-Hz response in the 0.25-s echoic condition (*p* = 0.4331, bootstrap, FDR corrected). When comparing with the anechoic condition, the 0.125-s echoic condition showed a significant reduction in phase coherence at the echo-related frequency (Figure 3A, *p* = 0.014 at 4 Hz, bootstrap, FDR corrected). The 0.25-s echoic condition showed a significant reduction at 6 Hz (*p* = 0.0005, bootstrap, FDR corrected) but not 2 Hz (*p =* 0.5879, bootstrap, FDR corrected). The reduction in 6-Hz phase coherence, however, was also observed in the 0.125-s echoic condition (*p* = 0.007, bootstrap, FDR corrected), which was not predicted by the neural adaptation models.

**Figure 4:**
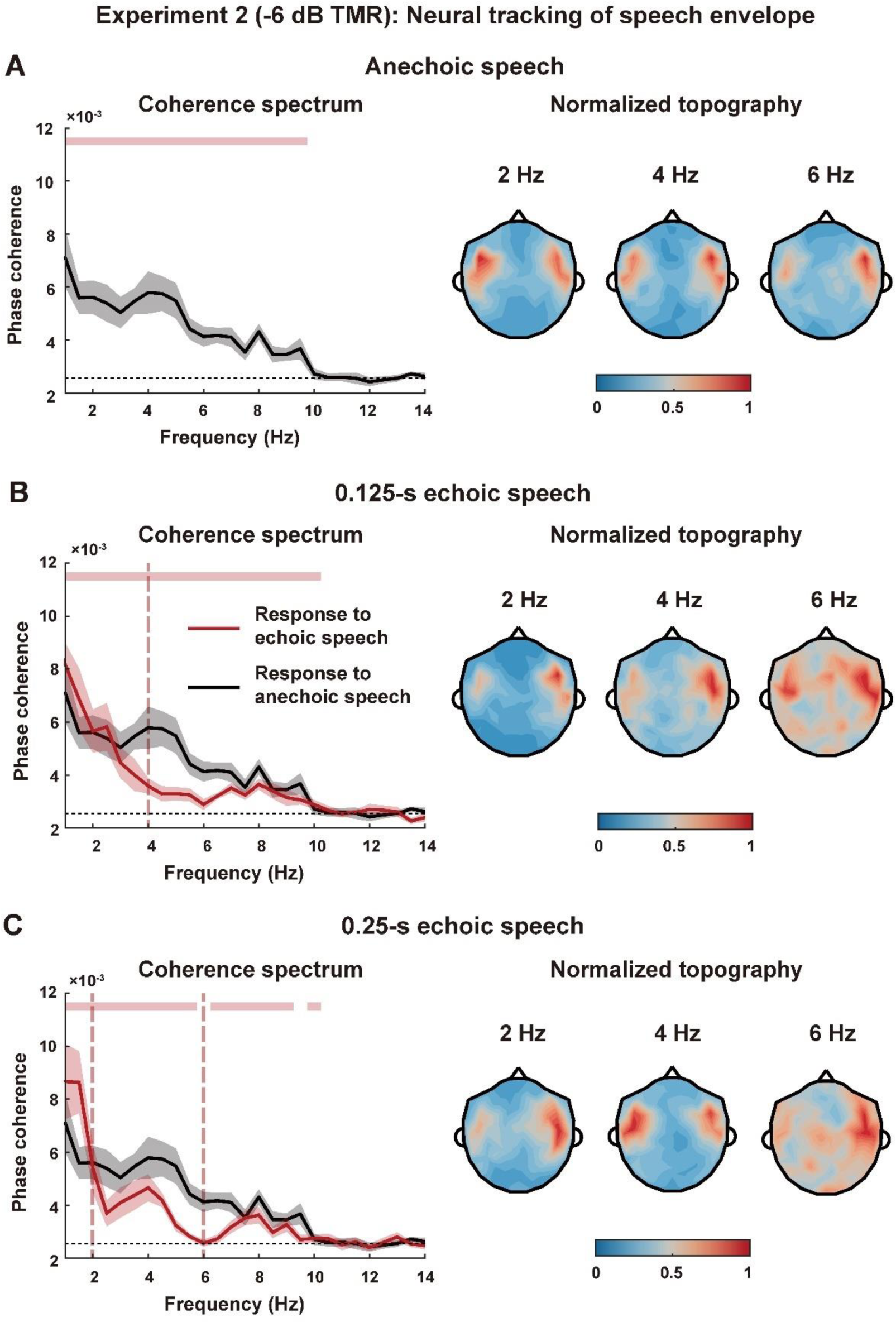
Phase coherence results of Experiment 2. Panels **A**, **B**, and **C** show the results in the anechoic condition, 0.125-s echoic condition, and 0.25-s echoic condition, respectively, with the same conventions as in Figure 2. **Left:** the phase coherence spectrum. **Right:** Topography of the phase coherence at echo-related frequencies.

### Neural segregation of direct sound and echo

The results of Experiments 1 and 2 suggested that cortical tracking of the speech envelope was insensitive to the presence of an echo, a phenomenon that could not be well explained by basic neural adaptation functions. Next, we further hypothesized that auditory stream segregation contributed to reliable neural tracking of speech envelope at echo-related frequencies. If echoic speech is neurally segregated into a direct sound and an echo, the two streams could be encoded by distinct spatio-temporal neural codes and described by a two-stream TRF model^12, 36^ (Figure 5A). We tested whether the streaming model could better explain the MEG response than a baseline model that modeled the MEG response using the temporal envelope of echoic speech (Figure 5A). We also compared the streaming model with an idealized model that modeled the MEG response only using the temporal envelope of direct sound (Figure 5A).

**Figure 5:**
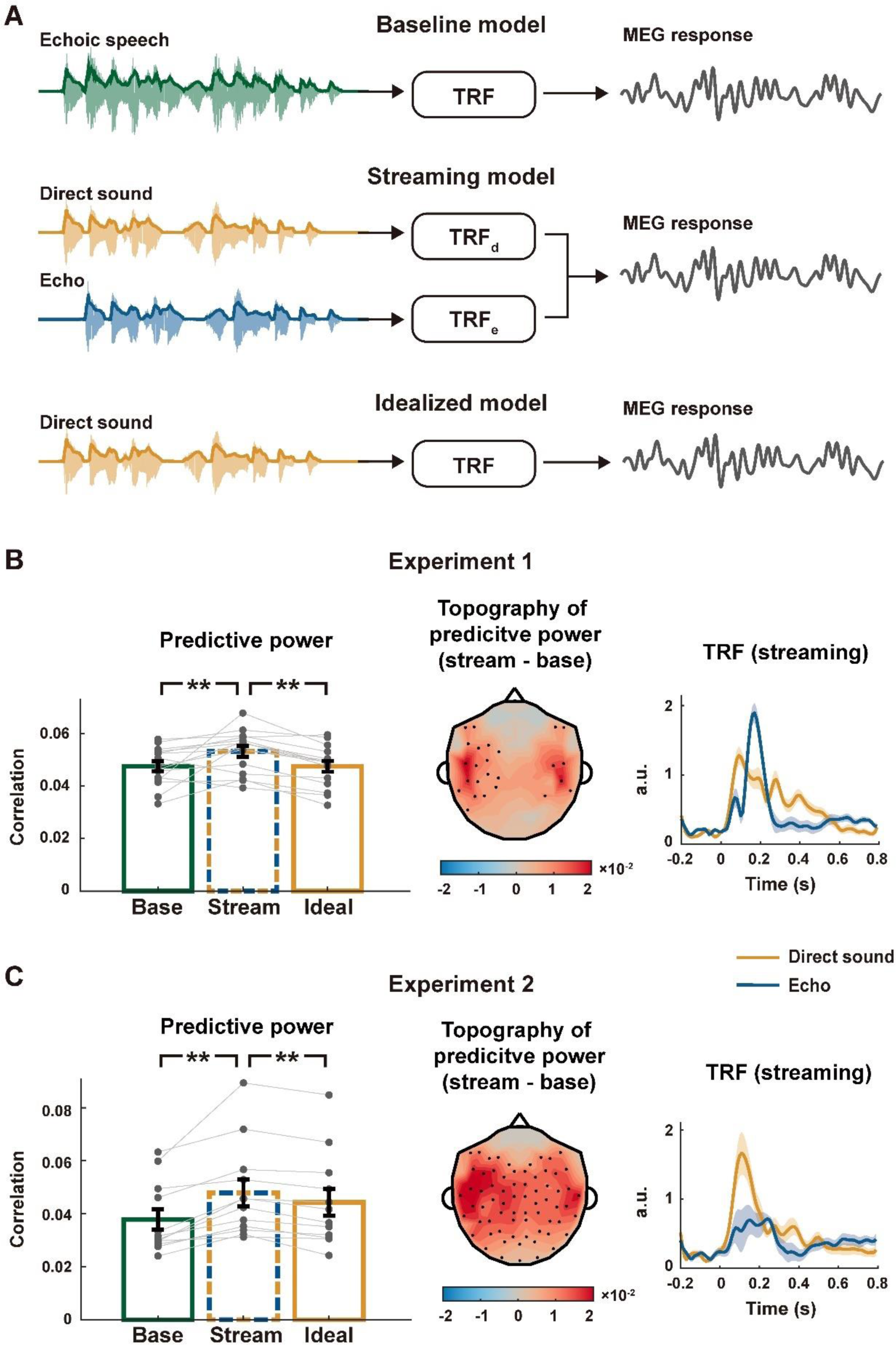
TRF model and results for Experiments 1 and 2. **A,** Illustration of three TRF models. The baseline model only considers the envelope of echoic speech, while the streaming model decomposes echoic speech into a direct sound and an echo and separately models their responses. The idealized model only considers the envelope of the direct sound. **B and C,** TRF results of Experiments 1 and 2. **Left:** The bars show the predictive power averaged over participants and MEG gradiometers, and the predictive power is significantly higher for the streaming model than the other two models (** *p* < 0.01, bootstrap, FDR corrected). Grey dots show the predictive power of individual participants. Error bars represent 1 standard error across participants (Experiment 1: *N* = 15; Experiment 2: *N* = 12). **Middle:** Topography of the difference in predictive power between the streaming model and the baseline model. The black dots indicate the sensor locations showing a significant difference between models (*p* < 0.05, bootstrap, FDR corrected). **Right:** TRF for the streaming model (averaged over participants and MEG gradiometers). Shaded areas cover 1 standard error across participants on each side.

In each echoic condition, the direct sound and echo were fully correlated except for a time delay. Therefore, to dissociate the neural responses to the direct sound and the echo, we pooled stimulus conditions with different echo delays (i.e., 0.125-s and 0.25-s delay) in the TRF analysis. The correlation coefficient between the actual MEG response and the response predicted by a TRF model was referred to as the predictive power. The predictive power averaged over all MEG gradiometers was shown in Figure 5B, C. In both experiments, the streaming model had higher predictive power than the baseline model (Experiment 1: *p =* 0.0012; Experiment 2: *p* = 0.0002, bootstrap, FDR corrected) and the idealized model (Experiment 1: *p =* 0.0004; Experiment 2: *p* = 0.0002, bootstrap, FDR corrected). When individual MEG channels were considered, the streaming model outperformed the baseline model mostly in the left hemisphere channels (Figure 5B, C). The TRF for the two streams were illustrated and the TRF to the direct sound tended to last longer (Figure 5B, C).

The TRF analysis suggested that the auditory system segregated the direct sound and the echo into separate streams. Next, we constructed additional conditions to further validate the conclusion. First, to further reduce the correlation between the direct sound and echo, we constructed a variable-delay echoic speech condition, in which the echo delay changed about every 3 seconds between 0.125 s and 0.25 s (Figure 6A). The variable-delay echoic speech also posed challenges for mechanisms that required a long adaptation time, e.g., adaptive filtering, but should barely affect auditory stream segregation. Second, speech segregation was enabled by speech segregation cues^3^. Therefore, if we removed the primary speech segregation cues, i.e., the spectro-temporal fine structure^19, 20^, neural segregation of speech should fail, but neural adaptation mechanisms should barely be influenced. Here, we removed the fine structure through 1-channel noise vocoding, which rendered speech unintelligible. Finally, we also manipulated the task to test whether top-down attention was required to segregate the echo from direct sound.

**Figure 6:**
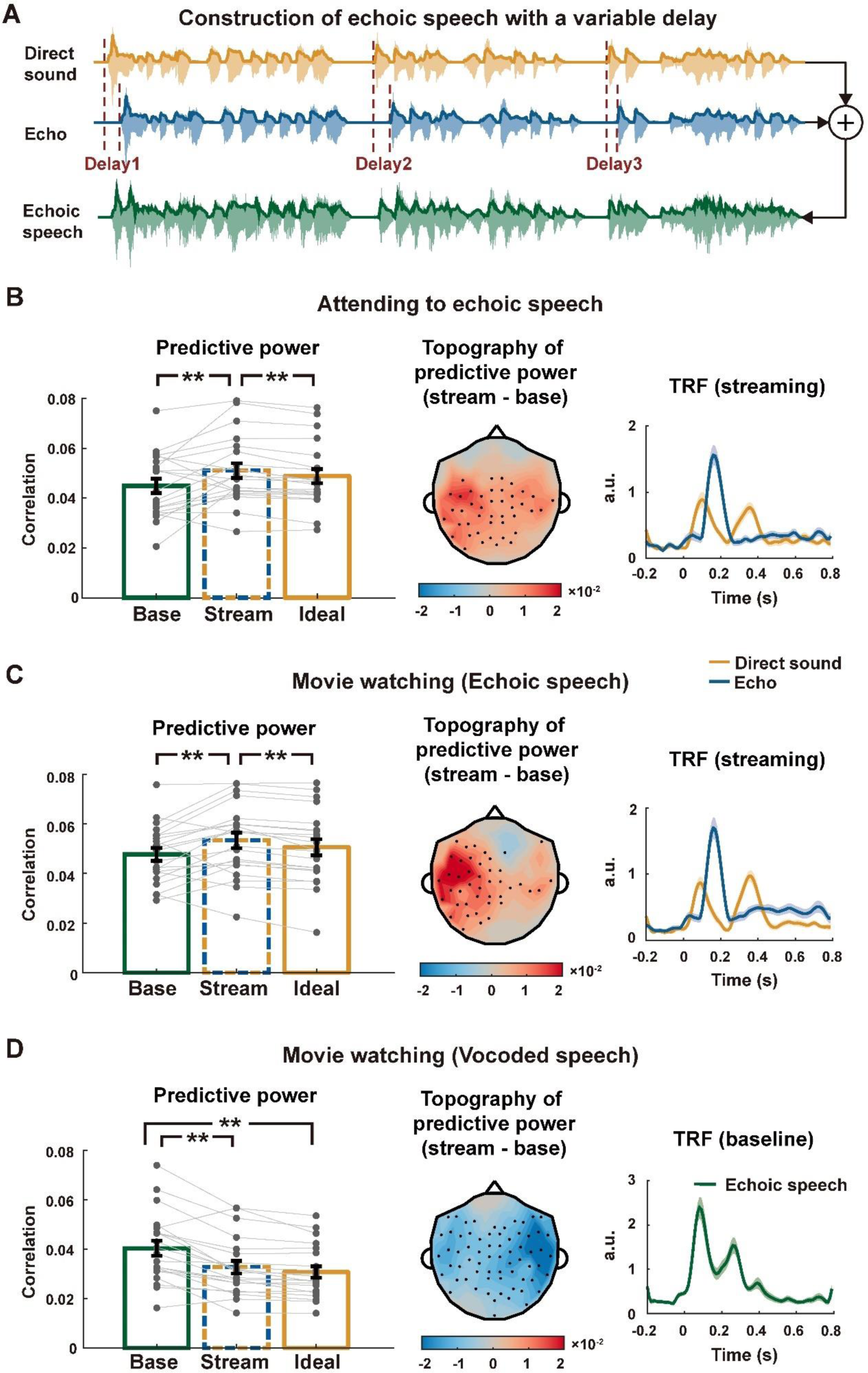
Results of Experiment 3. **A,** Construction of echoic speech with a variable delay. The echo delay changes about every 3 seconds and each time the delay is independently drawn from a uniform distribution between 0.125 s and 0.25 s. **B, C, and D,** TRF results. **Left:** Predictive power of three models (averaged over participants and MEG gradiometers). In panels **B** and **C**, the predictive power is significantly higher for the streaming model than the other two models. In panel **D**, the predictive power is significantly higher for the baseline model than the other two models (** *p* < 0.01, bootstrap, FDR corrected). Error bars represent 1 standard error across participants (*N* = 21). Grey dots show the predictive power of individual participants. **Middle:** Topography of the difference in predictive power between the streaming model and the baseline model. The black dots indicate the sensor locations showing a significant difference between models (*p* < 0.05, bootstrap, FDR corrected). **Right:** TRF of the model that best fits the neural response (averaged over participants and MEG gradiometers). Shaded areas cover 1 standard error across participants on each side.

Experiment 3 consisted of three conditions. In the first condition, the participants attended to variable-delay echoic speech and answered comprehension questions (accuracy = 95.2%). The second condition was the same as the first condition except that the participants watched a silent movie and were asked to ignore the echoic speech. In both conditions, the streaming model had higher predictive power than the baseline model (Figure 6B, C, left plot, *p* = 0.0058 and *p* = 0.003 for the 1^st^ and 2^nd^ conditions respectively, bootstrap, FDR corrected) and the idealized model (Figure 6B, C, left plot, *p* = 0.0004 for the 1^st^ and 2^nd^ conditions, bootstrap, FDR corrected). The advantage of streaming model lateralized to the left hemisphere channels (Figure 6B, C, middle plot). The third condition presented 1-channel noise vocoded echoic speech and the participants watched a silent movie while listening. In this condition, the baseline model better explained the MEG response than the streaming model (Figure 6D, left plot, *p* = 0.0002, bootstrap, FDR corrected) and the idealized model (Figure 6D, left plot, *p* = 0.0002, bootstrap, FDR corrected).

## Discussion

An intense echo strongly distorts speech envelope that contains critical cues for speech intelligibility, but human listeners can still reliably recognize echoic speech. The current study shows that the auditory system can effectively restore the low-frequency components of speech envelope that are attenuated or eliminated by an echo, providing a plausible neural basis for reliable speech recognition. Critically, for the conditions tested in the current study, the echo-related influence on speech envelope cannot be effectively compensated by basic neural adaptation mechanisms (Figure 1-Figure 4). Instead, the results strongly suggest that the auditory system pre-attentively segregates the direct sound and echo into different streams to maintain a faithful representation of speech (Figure 5 and Figure 6). These results show that auditory stream segregation is not just a key mechanism to achieve robust speech representation in multi-talker environments but may also play a critical role in separating repeated speech streams from the same talker.

Ecologically, an echo can be produced when a distant hard surface generates intense acoustic reflection with >50-ms delay^37^. Reflections with shorter delay are perceptually integrated with the direct sound and can benefit, instead of harming, speech recognition in noisy environments^38^. Echoes are perceptually salient acoustic phenomena. For example, it has been suggested that ancient rock art is often created at places that can generate maximal echo intensities^39^. During modern teleconferencing, echoes are frequently encountered: A echo is generated when the voice from one side of the conversation is picked up by the microphone on another side and transmitted back^31^. In real teleconference recordings, echoes almost always have >100 ms latency^40^. Although such long-latency echoes are not prevalent in natural environments, they do not diminish speech recognition, suggesting that the effects of echoes can be neurally compensated through existing mechanisms.

Neural adaptation is an obvious candidate to suppress the neural response to echoes, and previous studies have shown that neural adaptation to sound statistics can well suppress the influence of reverberation on speech encoding^9, 32^. Furthermore, when two brief sounds are presented within a few seconds, neural adaptation attenuate the response to the second sound^41^. Attenuating the echo to a long-lasting dynamic sound, such as connected speech, however, is much more challenging for at least two reasons. First, identifying which sound is an echo is a nontrivial question. For a brief sound, an echo can be defined as the lagging sound. In connected speech, however, a speech segment always follows another speech segment even without any echo, and therefore it is challenging to distinguish which segment belongs to the direct sound and which belongs to the echo. Second, since the direct sound and the echo overlap in time and frequency, a simple adaptation strategy cannot selectively suppress the echo while preserving the direct sound. Therefore, although neural adaptation can help to suppress echoes to some extent, it cannot well explain the robust cortical response to echoic speech. Furthermore, when the echo is more intense than the direct sound, synaptic depression tends to aggravate the effect of echo instead of canceling it (Figure 1D).

In highly complex listening environments, such as multi-speaker environments, auditory stream segregation is the basis for robust speech recognition^3, 4^. In such environments, different speakers generally differ in their pitch and spatial locations, which serve as bottom-up cues to segregate speech streams. When two utterances of the same speaker are mixed, they can be barely segregated based on bottom-up sensory features^5^. Nevertheless, when prior information is available, e.g., when part of the speech content is known, the brain can also use the prior information to separate speech streams, an effect referred to as schema-based speech segregation^2, 42^. At the neurophysiological level, it has also been shown that cortical activity can separately encode two speech streams produced by the same speaker when prior information is available^43^. For echoic speech, it is possible that the direct sound serves as prior knowledge to identify the echo. Furthermore, since speech is a dynamic sound, at each moment, the direct sound and echo differ in their pitch and spectral modulations, which provide cues to segregate the direct sound and echo into different streams. In natural listening environments, the direct sound and echo differ in their source locations, which can provide additional cues for speech segregation. Here, we do not investigate how speech segregation is achieved and potential mechanisms include, e.g., spectro-temporal masking^18^ and feature grouping based on temporal coherence^3^.

Here, we demonstrate that neural activity differentially encodes the direct sound and the echo, and the streaming model can better explain the MEG response than the baseline model (Figure 5 and Figure 6), providing evidence that auditory stream segregation underlies reliable perception of echoic speech. Previous studies have also suggested that the echo to a sound is also perceived and neurally encoded as an auditory object that is separated from the direct sound^33^. Furthermore, we demonstrate that a streaming model that considers both the direct sound and the echo can better explain the MEG response to echoic speech than a model that only considers the direct sound, consistent with recent studies showing cortical activity encodes properties of reverberant speech in addition to the direct sound^44, 45^.

Finally, the MEG response is better explained by the streaming model than the baseline model even when the listener’s attention is diverted by a silent movie (Experiment 3). Movie watching can attract attention but does not impose high processing demand. Therefore, the current results suggest that the segregation of direct sound and echo occurs with minimal involvement of top-down attention, and further studies may investigate whether the result remains when participants are distracted by more demanding auditory tasks. In the current study, the predictive power of the streaming model does not significantly differ between the story listening task and the movie watching task, consistent with previous results that the envelope-tracking response is minimally modulated by cross-modal attention^46^. It has been heavily debated whether auditory stream segregation requires attention. Some theories and empirical evidences suggest that top-down attention is necessary to parse a complex auditory scene into auditory streams^3, 47^, while others argue that primitive auditory streaming can occur preattentively^11, 48^. Auditory streaming is a complex process and it is possible that different modules involved in auditory streaming are differentially modulated by attention^44, 49, 50^, perceptual similarity between streams^51^, and task demand^52^. In the current experiment, the immediate repetition between direct sound and echo can serve as an effective cue for speech segregation, and previous studies have indeed demonstrated that auditory streaming based on repetitions in the sound pattern can occur without the engagement of top-down attention^53^.

In summary, the current study demonstrates that echo cancellation cannot be solved by neural adaptation mechanisms that can effectively cancel reverberation in typical daily acoustic environments. Instead, the processing of echoic speech involves speech segregation mechanisms that are more similar to the mechanisms engaged in processing speech in multi-talker environments. In other words, speech processing in an echoic environment is better viewed as an informational masking problem instead of an energetic masking problem^6^. These results provide a plausible explanation of why human listeners can tolerate echoes during online conferences and provide new insights into how to build brain-inspired echo cancellation algorithms: Although echoes and reverberations are both generated by acoustic reflections, echoes may be better suppressed using knowledge-guided speech segregation than algorithms optimized to cancel reverberations.

## Methods

### Participants

Totally, fifty-one participants (19∼33 years old, mean age, 24.6 years) participated in the study. Fifteen (5 male) participated in Experiment 1, Twelve (3 male) participated in Experiment 2, and twenty-four (11 male) participated in Experiment 3. Three participants of Experiment 3 were excluded since the trigger signaling sound onset was missing. All participants were right-handed native Chinese speakers, with no self-reported hearing loss or neurological disorders. The experimental procedures were approved by the Ethics Committee of the College of Biomedical Engineering & Instrument, Zhejiang University (No. 2022-001). The participants provided written consent and were paid.

### Stimuli

The speech stimulus was taken from the beginning of a narration of the story *Thatched Memories* by Wenxuan Cao. Three segments of recording were used in the experiment, each of which lasted about 13 minutes, with the constraint that the segment ended at the end of a sentence. The actual duration of the three segments was 12’59’’, 13’11’’, and 12’56’’, respectively. Each segment was used to generate the stimulus in one stimulus condition. Since each experiment consisted of three stimulus conditions, all three segments were presented in each experiment and their presentation order was kept the same, consistent with their order in the story. The order of stimulus conditions, i.e., how speech was manipulated, however was randomized and described in Procedures.

Echoic speech was constructed by delaying a speech signal by time *τ* and adding it back to the original signal (Figure 1A). If a speech signal was denoted as *s*(*t*), echoic speech could be expressed as *As*(*t*) + *s*(*t - τ*). The amplitude parameter *A* was 1 in Experiments 1 and 3, and was 0.5 in Experiment 2. When *A* was 1, the echo had maximal influence on the modulation spectrum and could completely notch out the speech envelope at echo-related frequencies. When *A* was 0.5, the echo was more intensive than the direct sound and simulations showed that neural adaptation slightly attenuated the neural response to the echo and rendered the neural responses to the echo and direct sound of more similar amplitude. Consequently, neural adaptation aggregated the influence of an echo on neural tracking of speech envelope at echo-related frequencies.

In Experiments 1 and 2, the delay *τ* was either 0.125 s or 0.25 s in a stimulus condition, and remained unchanged throughout each ∼13-min stimulus. We tested these two delays since a recent study demonstrated that a 0.125- and 0.25-s echo selectively attenuate the speech envelope at 4 Hz and 2 Hz^29^, respectively, and these modulation frequencies were critical for speech intelligibility^22, 25, 26^. In Experiment 3, the echoic speech had a variable delay. The delay varied whenever there was a pause longer than 500-ms in speech, and each time the delay was independently drawn from a uniform distribution between 0.125 s and 0.25 s. In the current speech materials, the interval between pauses longer than 500-ms was between 0.4-s and 11.2-s (3.4-s on average).

The 1-channel vocoded speech in Experiment 3 was constructed based on echoic speech with a variable delay. The speech envelope was extracted using the Hilbert Transform and was lowpass filtered below 160 Hz (4^th^-order Butterworth filter). The 1-channel vocoded speech was generated by modulating Gaussian white noise (filtered below 4 kHz) with the temporal envelope of echoic speech. The RMS intensity of the noise-vocoded stimulus was adjusted to match that of echoic speech without noise vocoding.

### Procedures

Participants sat in a quiet room while listening to speech, and their neural responses were recording using MEG. All speech stimuli were diotically presented through headphones. Each experiment lasted about 50 min.

#### Experiment 1

Experiment 1 consisted of three conditions that separately presented anechoic speech, echoic speech with a constant 0.125-s delay, and echoic speech with a constant 0.25-s delay. Each condition was presented in a separate block, and the order of the conditions was randomized among participants. While listening to speech, the participants fixated at the cross to reduce eye movements. The participants were instructed to attend to speech and answer two comprehension questions after each block.

#### Experiment 2

The procedure of Experiment 2 was identical to the procedure in Experiment 1, but the echoic speech differed in the TMR.

#### Experiment 3

Experiment 3 also consisted of three conditions, which presented echoic speech with a variable delay and vocoded echoic speech. The three conditions were presented in two blocks, the order which was randomized across participants. The first condition was presented in one block and the other two conditions were presented in the second block. The first condition presented echoic speech, and the participants performed the same question-answering task in Experiments 1 and 2. The second and the third conditions presented echoic speech and vocoded echoic speech, respectively. The two conditions were presented in the same block but their order was randomized among participants. In the block, the participants were asked to attend to a silent movie with subtitles (*The Little Prince*) and ignore any sound. Speech was presented about 5-min after the onset of movie, to make sure that participants were already engaged in the movie-watching task. No break was made between the two conditions with the same block, and the movie continued throughout. After the experiment, the participants had to answer simple comprehension questions about the movies and whether they noticed any auditory stimulus during the block. All participants answered the comprehension questions correctly.

### Data acquisition and preprocessing

Magnetoencephalographic (MEG) responses were recorded through a 306-sensor whole-head MEG system (MEGIN TRIUX^TM^ neo, Finland) at the Institute of Neuroscience, Chinese Academy of Sciences. The system had 102 magnetometers and 204 planar gradiometers, and the MEG signals were sampled at 1 kHz. All data preprocessing and analysis were performed using MATLAB (The MathWorks, Natick, MA). The temporal signal space separation (*tSSS*) was used to remove the external interference from the MEG responses^54^. Afterwards, the MEG signals were down-sampled to 100 Hz, and band-pass filtered between 0.8 and 10 Hz using a linear-phase finite impulse response (FIR) filter (6 s Hamming window, 6 dB attenuation at the cut-off frequencies). The linear-phase FIR filter caused a 3-s constant time delay to the input, which was compensated by removing the first 3-s samples in the filter output.

### Data analysis

#### Phase coherence spectrum

The phase coherence spectrum characterized the phase locking between the stimulus envelope and neural response, in each frequency band. The sound envelope was extracted by applying full-wave rectification to the sound, and low-pass filtering the signal below 50 Hz. The sound envelope and the neural response were both segmented into nonoverlapping 2-s time epochs, and each epoch was transformed into the frequency domain using the discrete Fourier transform (DFT). For each complex-valued DFT coefficient, the phase angle was extracted. The phase for stimulus envelope and neural response were denoted as *α_ft_* and *β_ft_* respectively, for frequency *f* and epoch *t*, and the phase difference between response and stimulus was therefore *θ_ft_* = *α_ft_* - *β_ft_*. The phase coherence spectrum was defined as follows:

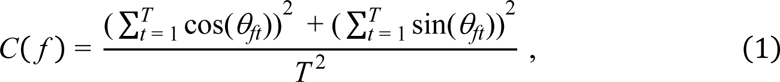

where *C*(*f*) was the phase coherence at frequency *f*, and *T* is the total number of epochs. The phase coherence was between 0 and 1. High phase coherence indicated that the phase difference between stimulus and response was consistent across epochs, i.e., stronger stimulus-response phase synchronization. The phase coherence spectrum was independently calculated for each MEG gradiometer.

#### Temporal response function

The temporal response function (TRF) was employed to model the time-domain relationship between sound envelope and MEG response. The baseline model assumed that the brain encoded the envelope of echoic speech, and the model was described as:

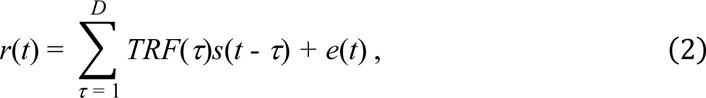

where *r*(*t*), *s*(*t*), and *e*(*t*) denoted the MEG response, envelope of echoic speech, and the residual error, respectively. *TRF*(*t*) was the TRF function, which could be interpreted as the response triggered by a unit power increase of the stimulus^16^.

A streaming model assumed that the brain could segregate the echo and the direct sound into two streams, and encoded the two streams in different manners. The model was formulated as

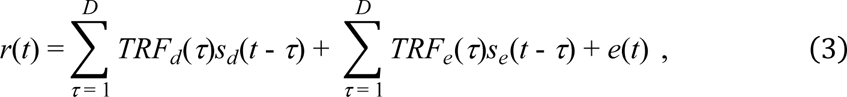

where *s_d_*(*t*) and *s_e_*(*t*) were the envelope of direct sound and echo respectively, and *TRF_d_*(*t*) and *TRF_e_*(*t*) were the TRF for direct sound and echo.

An ideal model assumed that the brain only responded to the direct sound and the model was formulated as

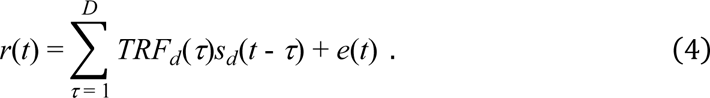

For all three models, the order of TRF, i.e., *D*, was 1 s, and the TRF is computed by performing the ridge regression with 10-fold cross-validation^55^. The predictive power of a model was defined as the correlation between a neural response and the TRF prediction. The regularization parameter for ridge regression was independently chosen for each model, to the maximize the predictive power in each experimental condition.

#### Response simulation using a synaptic depression model

We simulated how synaptic depression and gain control influenced envelope-tracking neural response. We adopted the model used by Mesgarani and colleagues, which could effectively reduce the influence of reverberation on speech responses^9^. The model input, i.e., presynaptic activity, was the auditory spectrogram, which contained 128 frequency channels and could be viewed as a simulation of subcortical auditory responses^56^. The auditory spectrogram was sampled at 200 Hz, and the temporal envelope in each frequency channel, referred to as *s*(*t*), was independently processed by the depression model. The neural response processed through the combined synaptic depression and gain control model was:

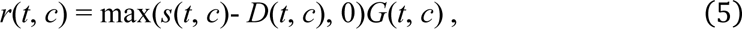

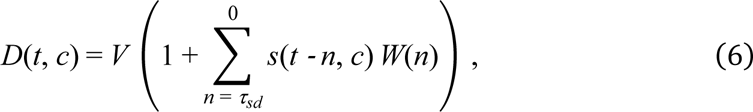

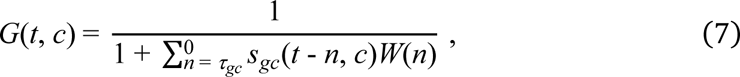

where *r*(*t*, *c*) was the model output, i.e., the simulated postsynaptic response. *W*(*n*) was a Hann window and the width equaled to *τ_sd_* and *τ_gc_* in *D*(*t*, *c*) and *G*(*t*, *c*) respectively. The simulated envelope-tracking response was obtained by summing the model output across all frequency channels. The model contained three parameters, i.e., *V*, *τ_sd_*, and *τ_gc_*, which were the firing threshold (0 < *V* < 1) and the time constants for synaptic depression and gain control (0 < *τ_sd_* < 500 ms, 0 < *τ_gc_* < 500 ms). We tested possible combinations of the parameters (step size = 0.05, 50 ms, and 50 ms, for *V*, *τ_sd_*, and *τ_gc_*) and chose the combination that could maximize the correlation coefficient between simulated neural response and the envelope of direct sound. The correlation coefficient was averaged over the 4 echoic conditions (i.e., 2 delay × 2 TMR). Another synaptic depression model^35^ and an optimal adaptive filter model^32^ were also simulated but they were less effective at restoring the speech envelope at echo-related frequencies (Figure S1).

### Statistical tests

To evaluate whether the phase coherence spectrum between stimulus and neural response at a frequency was significantly higher than the chance, the chance-level phase coherence spectrum was estimated using the following method^57, 58^. After the envelope of stimulus and neural response were segmented into 2-s time bins, we shuffled all time bins for the stimulus envelope so that the envelope and response were randomly paired. We calculated the coherence of the phase difference between randomly paired neural response and stimulus envelope. This procedure was repeated 5,000 times, creating 5,000 chance-level phase coherence values. Then the phase coherence values were averaged across channels and participants, for both the actual phase coherence and 5,000 chance-level phase coherence values. The significance level of the phase coherence at a frequency was (*M* + 1)/5,001 if it was lower than *M* out of the 5,000 chance-level phase coherence values at that frequency (one-sided comparison).

We used bias-corrected and accelerated bootstrap to compare the phase coherence value across conditions^59^. For two-sided paired comparisons, the data of all participants were resampled 10,000 times with replacement. If the resampled mean in one condition was greater than the resampled mean in the other condition for *M* out of the 10,000 replacements, the significance level was 2min(*M* + 1, 10,001 − *M*)/10,001. For two-sided unpaired comparisons, which was only used for the comparison across experiments, data in one condition were resampled 10,000 times. If the mean in the other condition was greater than *M* out of the 10,000 resampled mean values, the significance level was 2min(*M* + 1, 10,001 − *M*)/10,001. When multiple comparisons were performed, the *p*-value was adjusted using the false discovery rate (FDR) correction^60^.

## Data and code availability

The data and code to generate the figures can be found at: https://github.com/Yee-Gao/E-Encoding.

## Supplemental information

Supplemental methods and Figure S1.

## Acknowledgements

We thank the MEG facility of CAS CEBSIT/ION for the assistance in data collection, Wenyuan Yu and Jiajie Zou for constructive comments on an earlier version of the manuscript. This work was supported by the National Natural Science Foundation of China (32222035) and Key R & D Program of Zhejiang (2022C03011).

## Author contributions

N.D. conceived the study. N.D. and J.G. designed the experiment. J.G. and M.F. implemented the conducted the experiments. J.G. analyzed the data. N.D., J.G., and H.C. wrote the manuscript.

