## Supplemental method and Figure S1 for "Automatic Auditory Streaming Restores Missing Temporal Modulations in Echoic Speech"

**Supplemental information**

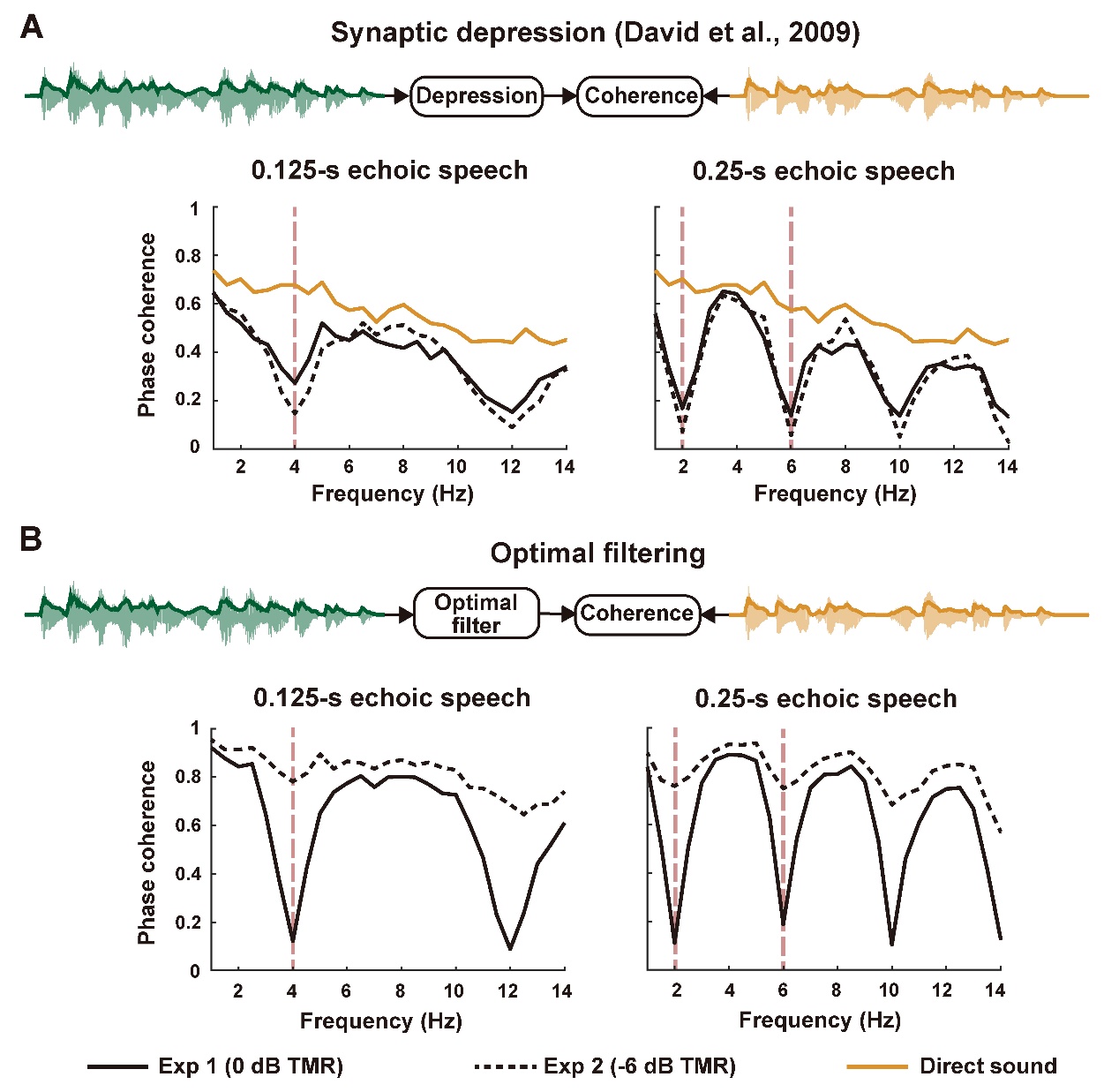

**Figure S1: Phase coherence spectrum between the simulated neural response to echoic speech and the envelope of direct sound.** **A,** Neural responses are simulated by processing the speech envelope using the synaptic depression model in David et al. (2009). **B,** Neural responses are simulated by processing the speech envelope using the optimal adaptive filter similar to that of Ivanov et al. (2022).

**Supplemental methods**

**Simulation of a synaptic depression model.**

The input to the synaptic depression model^1^, i.e., presynaptic activity, was the auditory spectrogram, which contained 128 frequency channels and each frequency channel was independently processed by the depression model. In the following, the level of depression was denoted as $\text{d }\text{(}\text{t}\text{, }\text{c}\text{)}$ for frequency channel *c* and time bin *t*. It was bounded between 0 and 1, and was initialized at 0 for *t* = 1. Its dynamics were described as:

$$\begin{aligned} \text{ }\text{d}\text{(}\text{t}\text{, }\text{c}\text{) }\text{= }\text{d}\text{(}\text{t }\text{- 1, }\text{c}\text{) }\text{+ }\text{s}\text{(}\text{t }\text{- 1, }\text{c}\text{)(1 - }\text{d}\text{(}\text{t }\text{- 1, }\text{c}\text{))}\text{v}\text{ - }\frac{\text{d}\text{(}\text{t}\text{- 1, }\text{c}\text{)}}{\text{τ}} \text{,}\#(1) \end{aligned}$$

where $\text{v}$ and $\text{τ}$ controlled the strength of depression and the time constant of recovery. To find the parameters that could best explain the neural response coherence spectrum, we scanned *v* in {0.05, 0.1, 0.2, 0.3, 0.4}/max(*s*(*t*, *c*)), and scanned $\text{τ}$ in {50, 100, 250, 500, 1,000, 1,500} ms. The postsynaptic response *r_d_* (*t*, *c*) was computed as follows:

$$\begin{aligned} \text{ }\text{r}\text{d}\text{(}\text{t}\text{, }\text{c}\text{) = }\text{y}\text{(}\text{t}\text{, }\text{c}\text{)(1 - }\text{d}\text{ (}\text{t}\text{, }\text{c}\text{))}\text{ }\text{,}\#\left( 2 \right) \end{aligned}$$

the simulated depressed neural response *r*(*t*), was obtained by summing the “depressed” spectrogram over frequency channels.

**Simulation of an optimal filter model.**

We simulated whether an optimal linear filter^2^ could effectively cancel the effects of echoes. The optimal filter *TRF_o_* was designed by solving the following equation using ridge regression.

$$\begin{aligned} {\text{ }\text{s}}_{\text{d}}\text{(}\text{t}\text{) }\text{=}\sum_{\text{τ}\text{ }\text{= 1}}^{\text{D}} \text{TRF}_{\text{o}}\text{(}\text{τ}\text{)}\text{s}\text{(}\text{t }\text{- }\text{τ}\text{) + }\text{e}\text{(}\text{t}\text{)} \text{,}\#\left( 3 \right) \end{aligned}$$

where *s*(*t*), *s_d_*(*t*), and *e*(*t*) denoted the envelope of echoic speech, the envelope of direct sound and the residual error, respectively. The output of the optimal filter, i.e., the simulated neural response, was

$$\begin{aligned} \text{ }\text{r}\text{(}\text{t}\text{) }\text{=}\sum_{\text{τ}\text{ }\text{= 1}}^{\text{D}} \text{TRF}_{\text{o}}\text{(}\text{τ}\text{)}\text{s}\text{(}\text{t }\text{- }\text{τ}\text{) .}\#\left( 4 \right) \end{aligned}$$

The optimal filter was not expected to completely remove the echo effect for the following reason. Adding an echo to a signal is equivalent to filtering the signal using a linear filter, and the transfer function of the linear filter corresponding to the echoic speech in the current study was *H*(*z*) = *A* + *z* ^–^ *^N^*, where *A* was the amplitude of the direct sound and *N* was the delay of echo in terms of the number of samples. In the current study, *A* was 1 or 0.5, rendering the transfer function irreversible. In other words, the echoes in the current study could not be completely cancelled by any linear filter.
